# Acute regulation of habituation learning via posttranslational palmitoylation

**DOI:** 10.1101/570044

**Authors:** Jessica C. Nelson, Eric Witze, Zhongming Ma, Francesca Ciocco, Abigaile Frerotte, J. Kevin Foskett, Michael Granato

**Affiliations:** Department of Cell and Developmental Biology; University of Pennsylvania, Perelman School of Medicine; 421 Curie Blvd, Philadelphia, PA, 19104; USA; Department of Cancer Biology; University of Pennsylvania, Perelman School of Medicine; 421 Curie Blvd, Philadelphia, PA, 19104; USA; Department of Physiology; University of Pennsylvania, Perelman School of Medicine; 415 Curie Blvd, Philadelphia, PA, 19104; USA; Department of Biology; Haverford College; 370 Lancaster Avenue, Haverford, PA 19041

## Abstract

Habituation is an adaptive learning process that enables animals to adjust innate behaviors to changes in the environment. Despite its well documented implications for a wide diversity of behaviors, the molecular and cellular basis of habituation learning is not well understood. Using whole genome sequencing of zebrafish mutants isolated in an unbiased genetic screen, we identified the palmitoyltransferase Hip14 as a critical regulator of habituation learning. We demonstrate that Hip14 regulates depression of sensory inputs onto an identified neuron and provide compelling evidence that Hip14 palmitoylates the Shaker-like channel subunit Kv1.1, thereby regulating Kv1.1 subcellular localization. Furthermore, we show that loss of either Kv1.1 or Hip14 leads to habituation deficits, and that Hip14 is dispensable in development and instead acts acutely to promote habituation. Combined, our results uncover a previously unappreciated role for acute post-translational palmitoylation at defined circuit components to regulate learning.

## Introduction

Habituation is an evolutionarily conserved non-associative form of learning characterized by a response decrement to repeated stimuli. One of the simplest forms of learning, habituation is defined by ten behavioral characteristics^1,2^, and has been associated with a wide range of behaviors and physiological responses including feeding and drug seeking^3^–^5^, neuroendocrine responses to stress^6^, and mechanosensory^7,8^, olfactory^9^ and acoustic startle responses^10^. Moreover, because habituation enables animals to focus selectively on relevant stimuli, it is thought to be a prerequisite for more complex forms of learning^2^. In fact, habituation is impaired in several human disorders that present with more complex cognitive and learning deficits including autism spectrum disorders, schizophrenia, attention deficit hyperactivity disorder, and Huntington’s disease^11^, suggesting that some of the molecular, genetic, and circuit mechanisms that act in the context of habituation extend well beyond this form of learning.

Over the past decade, significant progress has been made toward understanding the circuit mechanisms of habituation learning in both vertebrate and invertebrate systems. Studies on sensory habituation in *C. elegans* and *Aplysia*, and startle habituation in zebrafish and rodents have revealed that the synapses between sensory afferents and ‘command’ interneurons undergo plasticity during habituation^12^–^17^. Furthermore, genetic screens in *C. elegans* and *Drosophila* have been instrumental in identifying genes that regulate habituation learning^18^–^28^. Thus, while a picture of habituation in invertebrates is emerging, a comprehensive view of vertebrate habituation learning is largely absent. For example, how have the cellular and circuit mechanisms that guide habituation learning in the invertebrate nervous system been modified to regulate this process in the vertebrate nervous system? Similarly, does the increased complexity of the vertebrate nervous system dictate additional genes and molecular mechanisms to regulate habituation learning? To address these questions, we previously conducted an unbiased genome-wide genetic screen to identify the first set of mutants with deficits in vertebrate habituation learning^29^. Here, we report on the molecular identification of two of these mutants, which uncover a previously unappreciated role for posttranslational modification by the palmitoyltransferase Hip14 in regulating habituation, and identify a previously unknown substrate of Hip14, the Shaker-like K^+^ channel subunit Kv1.1. Moreover, we show that Hip14 regulates Kv1.1 protein localization to presynaptic terminals onto an identified neuron and demonstrate that Hip14 promotes habituation learning acutely. Combined, our results support a model by which Hip14-dependent palmitoylation ensures proper Kv1.1 localization to presynaptic sites critical for habituation learning and point to an emerging role for posttranslational palmitoylation in regulating behavioral plasticity.

## Results

### The palmitoyltransferase *hip14* regulates habituation learning *in vivo*

At five days of age, larval zebrafish exhibit robust habituation learning when confronted with a series of high intensity acoustic stimuli (25.9dB), as larvae rapidly learn to ignore these apparently innocuous stimuli and cease responding^30,31^. We have previously uncovered extensive pharmacological conservation of habituation learning between larval zebrafish and adult mammals^30^, and have recently shown that habituation is accompanied by increased inhibitory drive combined with decreased excitatory drive converging on an identified hindbrain neuron, the Mauthner (M) cell^14^. To identify genes that regulate habituation learning, we conducted a forward genetic screen using an assay for acoustic startle habituation^29^. In this assay, baseline startle responsiveness is first established by presenting larvae with 10 high intensity acoustic stimuli at a 20-second interstimulus interval (ISI) (**Fig. 1a**). To induce habituation learning, 30 additional stimuli are then presented with a 1 second ISI (**Fig. 1a**). Wild type animals exposed to this protocol rapidly cease responding to acoustic stimuli, exhibiting habituation rates of approximately 80%^29^ (**Fig. 1b**). In response to the same stimulation protocol, larvae carrying the *slow learner*^*p174*^ mutation exhibit almost no habituation learning (**Fig. 1b**).

**Figure 1.**
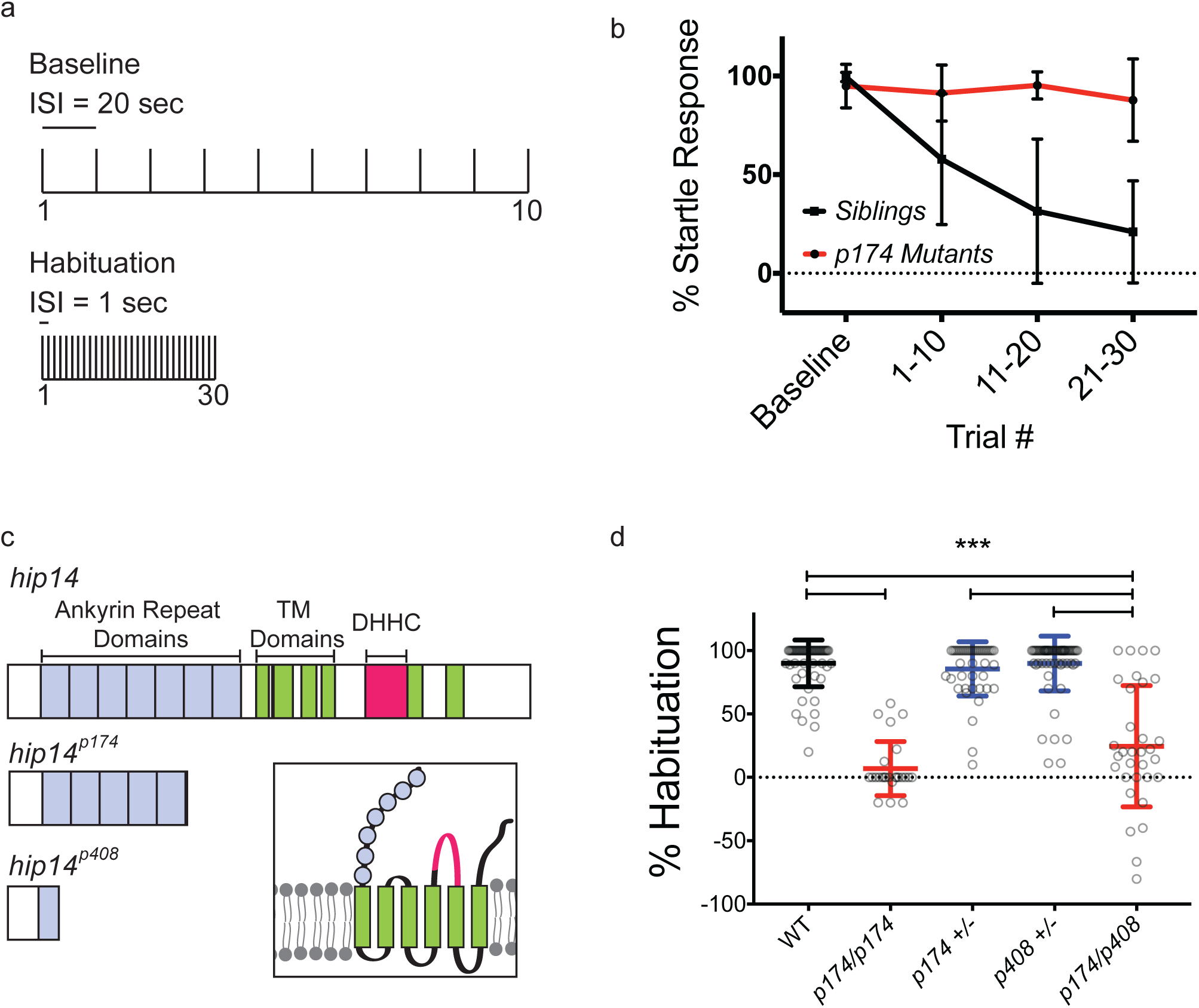
Hip14 regulates habituation learning in the larval zebrafish Hip14 is required for habituation learning in the larval zebrafish. (a) Acoustic stimulation protocol used to induce habituation learning. Vertical lines indicate acoustic pulses. 10 pulses are presented with 20-second ISI to assess baseline startle responsiveness. 30 pulses are then presented with 1-second ISI to induce habituation learning. (b) Habituation curves for sibling and *p174* mutant animals. Startle responses are averaged across bins of 10 stimuli. (Baseline= 10 stimuli at 20 second ISI, 1-10 = 1st bin of 10 stimuli at 1-second ISI, 11-20 = 2nd bin of 10 stimuli at 1-second ISI, 21-30 = 3rd bin of 10 stimuli at 1-second ISI). Mean response frequency within each bin ± SD are depicted, n≥19 larvae per genotype (c) Domain structure of Hip14, conceptual translations of *hip14* mutant alleles. Gray indicates ankyrin repeat domains, green indicates transmembrane domains, pink indicates DHHC catalytic domain. Inset depicts domain conformations. (d) Complementation testing to confirm mapping of habituation phenotype to *hip14* nonsense mutation. % Habituation = (1-[response frequency **21-30**]/[response frequency **baseline**])*100. Mean ± SD, n ≥25 larvae per genotype. *** indicates ANOVA with Tukey’s post-hoc test for multiple comparisons p<0.0001.

To identify the gene disrupted in *slow learner*^*p174*^ mutants, we took a previously described whole genome sequencing (WGS) approach followed by homozygosity mapping^29,32,33^. This uncovered a premature stop codon (Y211*) in the *huntingtin interacting protein 14* (*hip14*) gene, encoding a palmitoyltransferase (PAT) which catalyzes the addition of palmitate moieties to cysteine residues of substrate proteins^34^. Like other PAT family members, Hip14 contains transmembrane domains flanking a catalytic palmitoyltransferase (DHHC) domain^35^ (**Fig. 1c**). The premature stop codon in *slow learner*^*p174*^ mutants is located in the sixth of seven Ankyrin repeat domains, and thus precedes the catalytic palmitoyltransferase domain. To confirm that mutations in *hip14* indeed cause the observed habituation phenotype, we generated a second allele, *hip14*^*p408*^ using CRISPR-Cas9 genome editing techniques. This allele truncates the protein after amino acid 67 (**Fig. 1c**). We find that animals heterozygous for either *hip14*^*p408*^ or the original allele, *hip14*^*p174*^ show normal rates of habituation, while larvae carrying a combination of both mutant alleles (*hip14*^*p408/p174*^) fail to habituate at rates comparable to larvae homozygous for *hip14*^*p174*^ (NS, p=0.093). Together, these data establish that mutations in *hip14* cause habituation deficits (**Fig. 1d)**, demonstrating that the *hip14* palmitoyltransferase is required for habituation learning.

### Hip14 regulates synaptic depression of neurons of the acoustic startle circuit

The neuronal circuits that govern acoustic startle behavior in the larval zebrafish are well-defined and analogous to those described in other vertebrates^36^–^39^. Acoustic stimuli detected by hair cells within the inner ear are conveyed via the auditory (VIIIth) cranial nerve to the lateral dendrite of the startle command neuron (M-cell), which directly activates spinal motoneurons to execute a startle response (**Fig. 2a**)^39^–^41^. To examine whether Hip14 regulates habituation at the level of the Mauthner neuron, we performed Ca^+2^ imaging of Mauthner activity in *hip14* mutants. We have previously established an assay to simultaneously monitor M-cell Ca^+2^ transients and behavior (tail deflections)^14^ in head-fixed zebrafish larvae. Using this assay, we have shown that acoustic inputs above 12dB trigger Ca^2+^ signals that spread from the lateral dendrite to the M-cell soma to elicit tail deflections characteristic of the acoustic startle response (**Fig. 2b-d**)^14^. In wild type larvae repeated acoustic stimuli induce rapid habituation of acoustic startle behaviors (tail deflections), accompanied by reduced Mauthner Ca^+2^ signals **(Fig. 2d-e)**. In contrast, *hip14* mutants exposed to the same stimuli continue to display robust Ca^+2^ transients and their startle responses fail to habituate (**Figure 2d-e**). Moreover, unlike their sibling counterparts, *hip14* mutant dendrites exhibit little to no synaptic depression (**Fig. 2f**). Combined these results demonstrate that during habituation learning, Hip14 regulates synaptic depression at the level of the Mauthner lateral dendrite or upstream.

**Figure 2.**
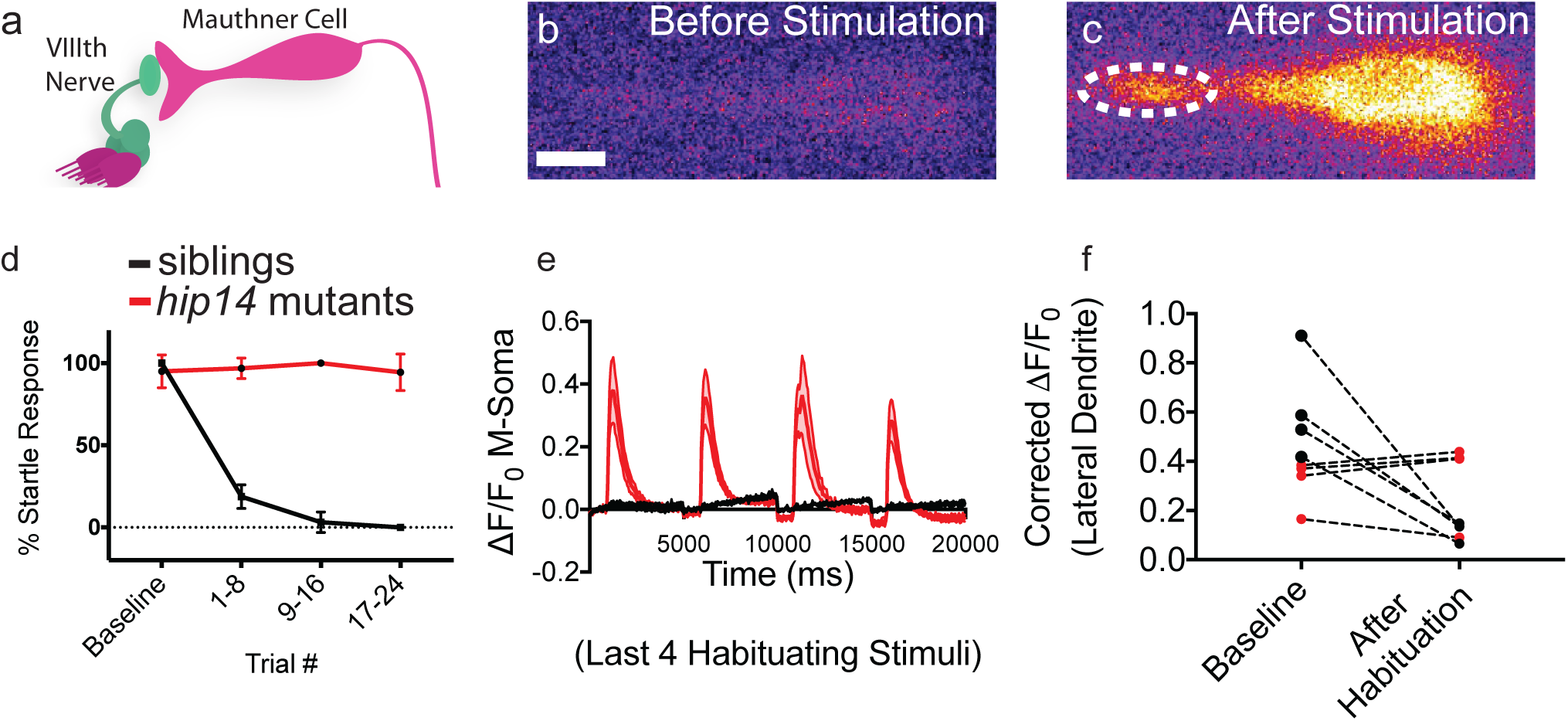
Lateral dendrite does not undergo synaptic depression in *hip14* mutants *hip14* mutants show no habituation at the level of the Mauthner lateral dendrite. (a) Diagram of auditory nerve inputs onto the Mauthner cell. The lateral dendrite, the site of these inputs, is one known locus for acoustic startle habituation. (b) Single frames showing GCaMP6s expressed in the Mauthner cell in an unstimulated animal and (c) following a high-intensity acoustic stimulus. (d) Habituation curves for head-embedded *hip14* mutant and sibling animals. Baseline = average of 5 responses at 2-minute ISI, 1-8 = 1st bin of 8 responses at 1-second ISI, 9-16 = 2nd bin of 8 responses at 1-second ISI, 17-24 = 3rd bin of 8 responses at 1-second ISI. Error bars indicate standard deviation. Mean ± SD, n=4 larvae per genotype. (e) Final 4 Mauthner soma responses (stimuli #21-24). By this point, sibling animals are fully habituated, and no longer performing startle responses. No Mauthner soma responses observed. *hip14* mutants show no/minimal behavioral habituation, and still have robust Mauthner responses. Mean ± shading indicates SEM, n=4 larvae per genotype. (f) Lateral dendrite responses before and after habituation in sibling and *hip14* mutant animals. Note the significant reduction in lateral dendrite Ca^+2^ activity in siblings (black), whereas *hip14* mutants show little/no such reduction (red) Mean lateral dendrite response, n=4 larvae per genotype. Unpaired t-test sibling lateral dendrite responses before as compared to after habituation, p=0.0152. Mutant lateral dendrite responses before as compared to after habituation, p=0.5308. Scale bar 10µm.

### Hip14 acts independently of previously identified regulators of habituation learning

To understand at the molecular level how *hip14* promotes habituation learning we wanted to identify the relevant Hip14 substrates. Hip14 was originally identified in the context of its interaction with the Huntington’s disease-associated Huntingtin protein^42^. Since its discovery, hundreds of potential Hip14 palmitoylation substrates and interacting partners have been identified^34,35,43^–^50^. Interestingly, multiple Hip14 interacting partners are components of pathways we previously implicated in habituation learning in the larval zebrafish. First, Hip14 interacts with PI3K and AKT activators that we previously demonstrated promote acoustic startle habituation downstream from the PAPP-AA / Insulin Growth Factor Receptor (IGF1R) signaling pathway (**Fig. 3a**)^29,43^. Second, Hip14 also interacts with PDE4, which acts downstream from *neurofibromatosis-1* (*nf1*) to regulate visual habituation learning in larval zebrafish (**Fig. 3a**)^43,51^. Importantly, habituation deficits in zebrafish *papp-aa* or *nf1* mutants can be restored by pharmacological manipulation of PI3K, AKT, cAMP and RAS signaling^29,51^. To determine whether these intracellular signaling pathways also function downstream of Hip14 to regulate habituation learning, we used the same pharmacological manipulations and found that manipulation of these signaling pathways in *hip14* mutants failed to restore normal habituation learning (**Fig. 3b-d**, **Supplemental Fig. 1)**. These data suggest that Hip14 acts independently of these previously identified learning pathways.

**Figure 3.**
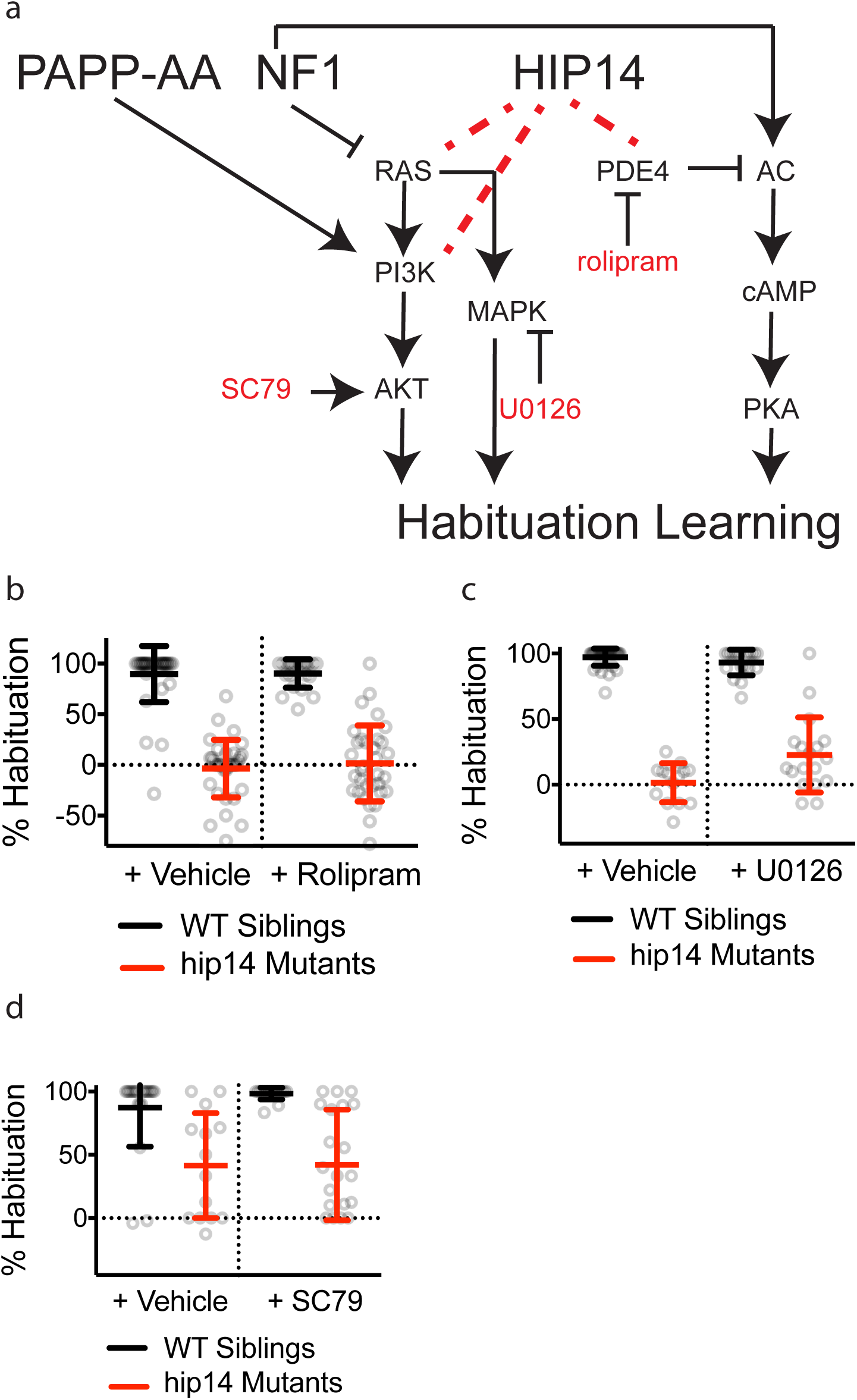
Hip14 regulates a novel learning pathway Hip14 does not act through previously established habituation pathways. (a) Pathway diagram indicating cellular mechanisms of action for PAPP-AA and NF1. Red text denotes drugs that impinge on individual pathway elements. Dotted lines indicate biochemical interactions between proteins that may or may not play a functional role in regulating habituation learning. (b-d) Habituation rates in *hip14* mutants and sibling larvae are unaffected by pharmacological manipulation of known habituation pathways: (b) 10μM Rolipram (or DMSO control) applied 30 minutes prior to and throughout behavior testing. (c) 1μM U0126 (or DMSO control) applied 30 minutes prior to and throughout behavior testing. (d) 1μM SC-79 (or DMSO control) applied from 3dpf-5dpf and throughout behavior testing.

### The Shaker channel K^+^ subunit Kv1.1 regulates habituation learning

Given the evidence that Hip14 acts independently of some of the known learning-relevant signaling pathways, we turned to our forward genetic screen, reasoning that other mutants with habituation deficits might represent potential Hip14 interacting partners. In particular, *fool-me-twice*^*p181*^ mutants show strong deficits in habituation learning similar to those observed in *hip14* mutants (**Fig. 4a**). Whole genome sequencing of *p181* mutants revealed a missense mutation leading to an amino acid substitution (N250K) in the coding sequence for *kcna1a*, which encodes the Shaker-like voltage-gated K^+^ channel subunit Kv1.1 (**Fig. 4b-c**). Shaker K^+^ channels are critical regulators of neuronal activity; in mutants lacking Shaker and Shaker-like K^+^ channels, neuronal repolarization is delayed^52^, neurons are hyperexcitable^53^ and release more quanta upon stimulation^54^, and animals display behavioral hyperexcitability and seizure-like behaviors^55,56^. Interestingly, previous studies have revealed that Kv1.1 requires palmitoylation for voltage sensing^57^ by an as-yet-unidentified PAT.

**Figure 4.**
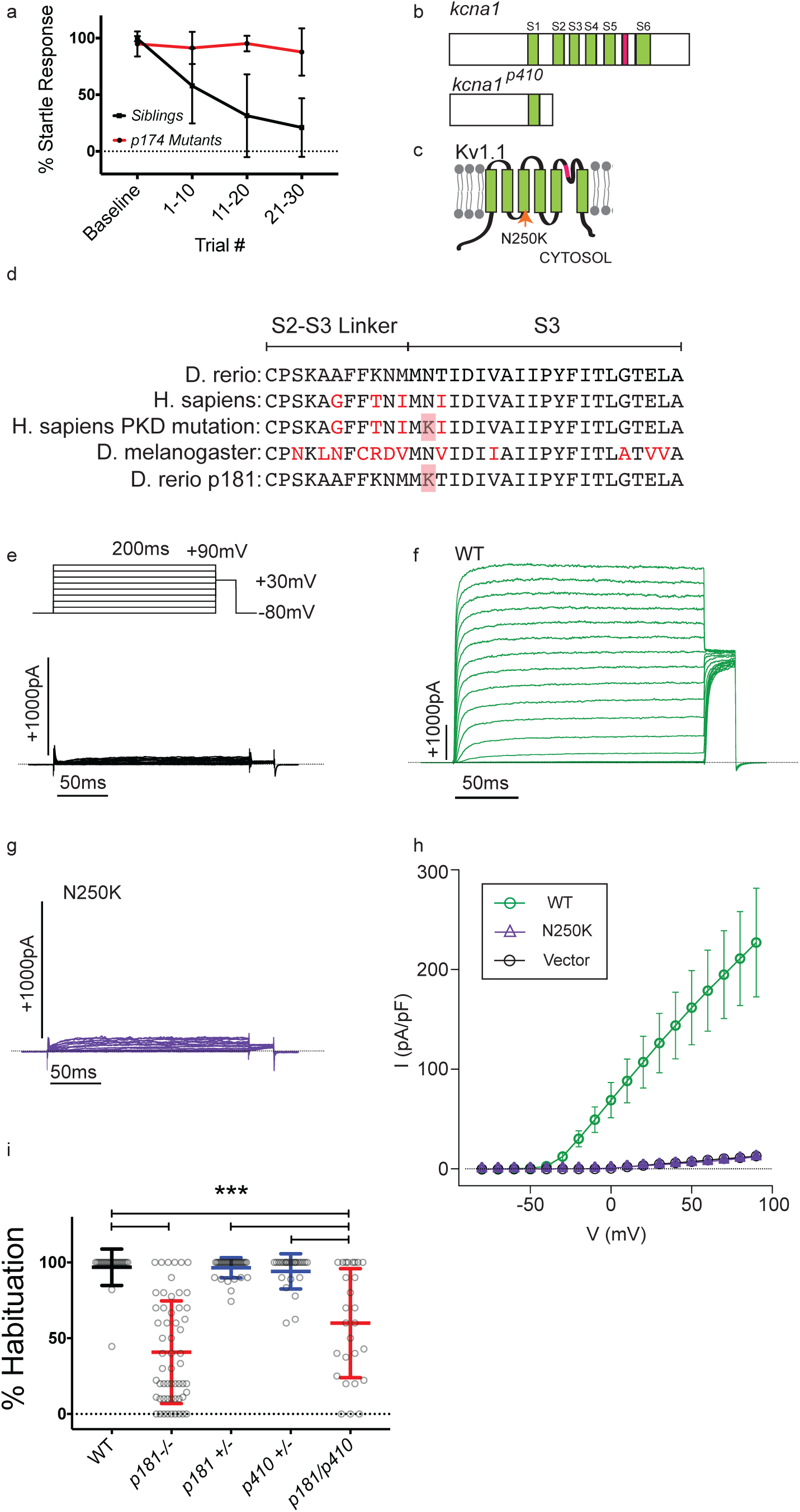
Mutations in *kcna1* disrupt habituation learning and result in a non-functional Shaker-like channel subunit Kv1.1 is required for habituation learning in the larval zebrafish. (a) Habituation curves for *p181* mutant animals and siblings. Startle responses are averaged across bins of 10 stimuli (as in **Fig. 1a**) Mean response frequency within each bin ± SD are depicted n≥57 per genotype (b) Conceptual translations of *kcna1a* mutant alleles, (c) domain structure of Kv1.1. Green indicates transmembrane domains, pink indicates pore helix domain. Orange arrow indicates N250K missense mutation identified in *p181*. (d) Alignment demonstrating strong conservation of Kv1.1 S2-S3 linker and S3 domain sequences. Note that the N250K missense mutation identified in our screen is equivalent to the N255K missense mutation associated with human PKD disease. **(e-g)** Representative whole-cell K^+^ family currents evoked by 200-ms voltage pulses from −80 to +90mV in 10mV increments from a holding potential of −80mV as in **Fig 1e**-top in N2a cells transfected with **(e)** vector only (control); cell capacitance of 15.5 pF, (**f**) wild type Kv1.1; cell capacitance of 18.2 pF, (**g**) KV1.1^N250K^; cell capacitance of 14.9 pF. For details of solutions see the Methods section. Dashed lines indicate 0-current level. (**h**) Steady-state current-voltage (I-V) relations for Vector (control), wild type Kv1.1 and KV1.1^N250K^, obtained by measurements of the currents at the end of 200-ms pulses, normalized by the cell capacitance. e.g., the normalized currents at +90 mV are 226.9 ± 54.5 (pA/pF) for wild type (n=6), 12.8 ± 1.0 (pA/pF) for KV1.1^N250K^ (n =7), and 11.2 ± 1.2 (pA/pF) for vector (n = 5). Two-tailed Student’s unpaired t-test: P=0.003, t_9_ = 3.89 for wild type and P = 0.315 and t_10_=1.06 for KV1.1^N250K^, compared to vector, indicating KV1.1^N250K^ is a nonfunctional channel. The cells used in these measurements had the whole-cell capacitance of 14.8 ± 1.6 (pF) (n = 5) for vector, 15.4 ± 0.8 (pF) (n = 6) for wild type, and 14.9 ± 1.3 (pF) (n = 7) for KV1.1^N250K^, respectively. (i) Complementation testing to confirm mapping of habituation phenotype to *kcna1a* missense mutation. Mean ± SD, n ≥23 larvae per genotype. *** indicates ANOVA with Tukey’s post hoc test for multiple comparisons p<0.0001.

The residue mutated in *fool-me-twice*^*p181*^ mutants, N250, is highly conserved across species, and this precise residue substitution is associated with human disease^58^, suggesting that the N250K missense mutation likely affects Kv1.1 function (**Fig. 4d**). To determine whether the Kv1.1^N250K^ substitution alters channel properties, we performed whole-cell current recordings in N2a cells expressing wild type and mutant versions of Kv1.1. This revealed that the Kv1.1^N250K^ substitution completely abolishes conductance, consistent with the hypothesis that the habituation deficit observed in *p181* mutant zebrafish is caused by a non-functional Kv1.1 (**Fig. 4e-h**). Finally, we used CRISPR-Cas9 genome editing to generate a zebrafish line carrying a premature stop codon in *kcna1a* (*p410*). Complementation analysis with the original *kcna1a*^*p181*^ allele confirmed that loss-of-function mutations in *kcna1a* cause deficits in habituation learning (**Fig. 4i**).

### Presynaptic Kv1.1 localization requires Hip14-dependent palmitoylation

Both Hip14 and Kv1.1 regulate habituation learning. Because Kv1.1 requires palmitoylation by an as-yet-unidentified PAT, we wondered whether Hip14 palmitoylates Kv1.1. To test this hypothesis, we co-expressed Flag-tagged recombinant Kv1.1 and HA-tagged recombinant Hip14 in HEK cells, and then performed an Acyl Biotin Exchange (ABE) assay^59^ to measure palmitoylation. We found that in the absence of Hip14, Kv1.1 is not palmitoylated above background levels (**Fig. 5a**). In contrast, co-expression of Kv1.1 with Hip14 resulted in robust Kv1.1 palmitoylation, demonstrating that Hip14 can palmitoylate Kv1.1 (**Fig. 5a**). To show that Hip14 catalytic activity is indeed required for Kv1.1 palmitoylation, we mutated the catalytic DHHC domain (hip14-DHHS). In contrast to co-expression with wild type Hip14, co-expression of Hip14-DHHS with Kv1.1 yielded dramatically reduced levels of palmitoylated Kv1.1 (**Fig. 5a**). Moreover, co-expression of wild type Hip14 with the Kv1.1^N250K^ mutant isolated in the genetic screen produced a marked reduction in palmitoylated Kv1.1 (**Fig. 5a**). Thus, Hip14 acts as a Kv1.1 palmitoyltransferase, consistent with the idea that Hip14 promotes habituation learning through Kv1.1.

**Figure 5.**
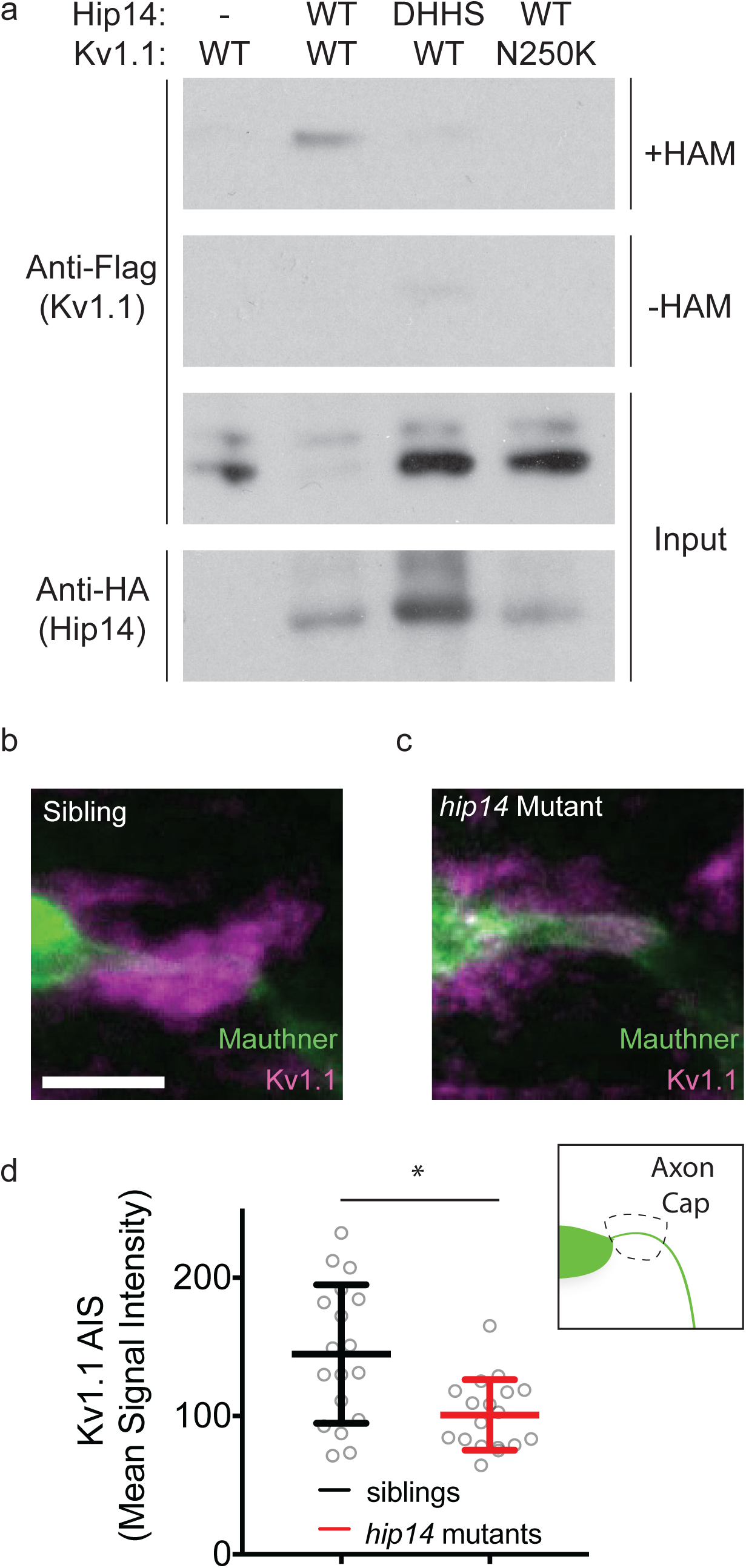
Hip14 palmitoylates Kv1.1, and regulates its localization in vivo. Hip14 palmitoylates Kv1.1 and regulates its localization *in vivo*. (a) Acyl Biotin Exchange assay, in which Kv1.1-Flag +/-Hip14-HA are transfected into HEK cells. Immunoblot indicates palmitoylation of Kv1.1-Flag in lanes with functional co-transfected Hip14-HA, whereas Kv1.1 is not palmitoylated in the absence of functional Hip14. Samples without hydroxylamine (-HAM) are negative controls for the ABE reactions. (b) Immunohistochemical labeling of the Mauthner cell (green) and Kv1.1 (magenta) in *gffDMC130a; uas:gap43:citrine* sibling, using Chicken anti-GFP, and Rabbit anti-Kv1.1 antibodies. Note the bright localization of Kv1.1 to the axon cap: synaptic inputs from spiral fiber neurons onto Mauthner cell AIS. (c) Immunohistochemical labeling of a *hip14* mutant animal as in (b). Note the reduction in Kv1.1 signal at the axon cap. (d) Quantification of Kv1.1 signal in mutants as compared to WT. Mean signal intensity = pixel intensity for axon cap region of interest (inset) divided by axon cap area. Unpaired t-test p=0.0021. Scale bar 10µm. Mean ± SD, n=18 axon caps per genotype (n≤2 axon caps per animal).

Post-translational palmitoylation is known to regulate substrate function as well as substrate localization^60^. Therefore, we hypothesized that Hip14-dependent palmitoylation might regulate Kv1.1 localization. To test this, we turned to immunohistochemistry and examined the localization of Kv1.1 in *hip14* mutant and sibling animals. Previous work in 3-day-old zebrafish larvae has shown that Kv1.1 is expressed throughout the hindbrain, including the Mauthner soma, and that *kcna1a* mRNA is present in the Mauthner cell as late as 5 days post-fertilization^61,62^. We find that at day 5, Kv1.1 protein expression persists throughout the CNS, including in hindbrain neurons of the startle circuit, yet only very weakly localizes to the Mauthner soma or lateral dendrite. Instead, Kv1.1 strongly localizes to a structure termed the axon cap (**Fig. 5b-d**), which is comprised of dense synaptic terminals from feed-forward spiral fiber neurons that wrap around the axon initial segment of the Mauthner axon^63,64^. Given the low Kv1.1 levels at the lateral dendrite and the strong accumulation of Kv1.1 in the axon cap, we selected the axon cap to determine if and to what extent Kv1.1 localization requires *hip14*. Analysis of *hip14* mutants revealed a significant reduction of Kv1.1 protein localization at the axon cap (**Fig. 5b-d**). Together, these biochemical and immunohistochemical results provide compelling evidence that Kv1.1 is a previously unrecognized Hip14 substrate, and that Hip14 regulates Kv1.1 localization to presynaptic terminals of the acoustic startle circuit. Combined, this supports a model by which neuronal Hip14 acts via Kv1.1 to promote habituation learning.

### Hip14 acts acutely to regulate habituation learning

Palmitoylation has previously been shown to function during development in the establishment of neural connectivity, as well as acutely in response to neural activity^60^. In neurons, palmitoylation can occur in both somatic Golgi, as well as in axons and dendrites^60^. Given that Hip14 has been observed in axons and presynaptic sites^47,65^, it is conceivable that Hip14 promotes habituation learning during development by helping to establish the habituation circuitry or acutely, for example at synapses by dynamically palmitoylating effector proteins. To distinguish between these possibilities, we employed an inducible transgenic rescue strategy to assess whether Hip14 expression after startle circuit development is sufficient to restore habituation learning. Development of the major components of the acoustic startle circuit is well-defined and largely occurs between 24 and 96 hours post-fertilization (hpf)^66^. Indeed, larvae perform acoustic startles and are capable of habituation learning at 96hpf indicating that the circuit elements required for habituation are in place by this time (**Fig. 6a**). We find that inducing transgenic Hip14 expression in otherwise *hip14* mutant animals as late as 3 hours prior to behavioral testing at 120hpf restored habituation learning (**Fig. 6c-d**). Thus, Hip14 acts after the development of startle circuit elements required for habituation, supporting a model by which Hip14-dependent palmitoylation and localization of Kv1.1. acutely promotes habituation learning.

**Figure 6.**
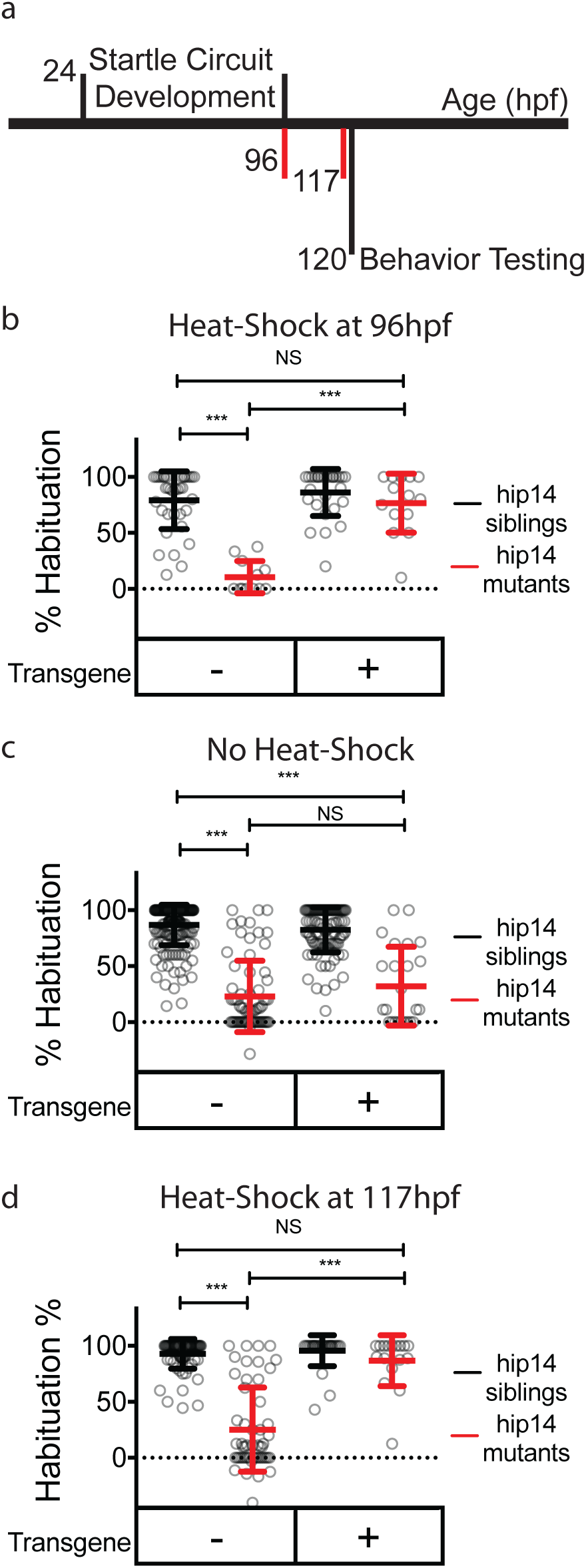
Hip14 rescues acutely Hip14 acutely regulates habituation learning. (a) Timeline depicting startle circuit development and heat shock timing. Numbers indicate hours post-fertilization. Red vertical lines indicate start times for heat-shock (96 hours in (b) and 117 hours in (d)) (b) Rates of habituation in animals heat-shocked at 37°C at 96 hours post-fertilization (4dpf). (c) Rates of habituation in animals without heat-shock. (d) Rates of habituation in animals heat-shocked at 37°C at 117 hours post-fertilization (5dpf). In b-d, testing is always performed at 120hpf (5dpf). Transgene = hsp70:hip14:p2a:mkate; Mean ± SD, n ≥12 per genotype; *** indicates p<0.0001; NS = Not Significant.

## Discussion

Habituation learning was initially described in the context of reflex responses^1,8^, however there has since been broad recognition that habituation learning is a widespread mechanism associated with and to some degree prerequisite for more complex cognitive processes and behaviors^3^–^6,11^. While our understanding of habituation learning both at the molecular and cellular level has been informed predominantly by molecular-genetic studies in invertebrates^18^–^28^, whether and how additional genetic and circuit mechanisms regulate vertebrate habituation learning has remained largely unclear. Here we describe a previously unappreciated role for the palmitoyltransferase Hip14 in regulating habituation learning, identify the Shaker-like channel subunit Kv1.1 as a Hip14 substrate that also regulates learning, and demonstrate that Hip14 acts to properly target Kv1.1 to presynaptic sites. Finally, we show that Hip14 acts acutely to promote habituation learning. Combined our results provide compelling evidence for Hip14-dependent palmitoylation as a dynamic regulator of habituation at a specific learning-relevant synapse.

Palmitoylation has received considerable attention as a post-translational modification regulating synaptic plasticity^60,67^–^71^. Furthermore, there is strong evidence that Hip14-dependent palmitoylation plays important roles in the nervous systems of invertebrates and vertebrates. For example, *Drosophila* Hip14 localizes to presynaptic terminals where it regulates their development^65^. In rodents *hip14* is implicated in a variety of behaviors^44,72,73^, yet the relevant substrates and whether *hip14* acts acutely or as a regulator of neuronal development or maintenance in these contexts has not been examined. Thus, our findings that Hip14 acts acutely during learning or just before to ‘prime’ learning-relevant synapses reveals a previously unappreciated dynamic role for Hip14-dependent palmitoylation. It is important to note that Hip14 is fairly unique among palmitoyltransferases in that its N-terminus is comprised of ankyrin-repeat domains that may confer additional functions beyond palmitoylation. This raises the intriguing possibility that Hip14 may act in part through palmitoylation-independent mechanisms to localize Kv1.1 protein, and future experiments are required to test this idea. Similarly, as palmitoylation is dynamically regulated, it will be interesting to determine whether dynamic regulation of protein de-palmitoylation is implicated in learning. Finally, while it is possible that additional palmitoyltransferases play critical roles in habituation learning, our finding that *hip14* is required for habituation and acts via a learning-relevant substrate, Kv1.1 supports the notion that palmitoylation is a highly selective process executed at selective sites by distinct palmitoyltransferases and their substrates.

Our results also suggest an evolutionarily conserved role for Kv1.1, a Shaker-like channel subunit identified in *Drosophila* for opposing roles in enhancing and suppressing habituation in different contexts^74,75^. Importantly, the identical amino acid substitution Kv1.1^N250K^ recovered in our genetic screen has previously been identified in humans (Kv1.1^N255K^) as associated with the movement disorder familial paroxysmal kinesigenic dyskinesia^58^. Previous work indicates that this mutation does not affect membrane expression of Kv1.1, but strongly reduces channel currents^58^, consistent with our finding that currents are abolished in cells expressing the Kv1.1^N250K^ allele. Here we show that this mutation also affects Hip14-dependent Kv1.1 palmitoylation. Given the importance of palmitoylation for Kv1.1 gating and function^57^, we propose that disrupted palmitoylation may represent one way in which this *kcna1* mutation leads to disrupted channel function and voltage gating. Indeed, multiple lines of evidence support our hypothesis that Hip14 regulates Kv1.1 localization and function through palmitoylation. First, mutants for each gene show a similar behavioral phenotype. Second, *in vitro* Hip14 palmitoylates Kv1.1, but fails to palmitoylate the learning-defective point mutant version of Kv1.1. Lastly, *in vivo* Hip14 is required for Kv1.1 protein localization to defined presynaptic sites, providing compelling evidence for a model by which dynamic Hip14-dependent palmitoylation of Kv1.1 influences its localization to defined presynaptic sites to promote learning.

Although it is viewed as a simple form of learning, previous work has suggested that multiple and potentially intersecting pathways drive habituation. We have previously shown that the M-cell lateral dendrite undergoes synaptic depression during habituation learning^14^; we now demonstrate that depression at the lateral dendrite is disrupted in Hip14 mutants, consistent with a role for Hip14 and Kv1.1 at this defined circuit locus. It is possible that to promote habituation learning Hip14 and Kv1.1 are additionally engaged at other neural circuit loci, including at the Mauthner cell axon cap where synaptic inputs from spiral fiber neurons terminate on the Mauthner AIS. Additional experiments are required to determine the precise cell type(s) and synapse(s) at which Hip14/Kv1.1 function to promote acoustic startle habituation in zebrafish. Similarly, it will be interesting to determine whether other molecular regulators of habituation identified in the zebrafish, such as Ap2s1-dependent endocytosis^29,32^; glutamate signaling^51^; NF1 signaling through cAMP and Ras^51^; or insulin growth factor signaling through PI3K and AKT^29^ function at the same or different circuit loci, and on the same or different time scales. Together with the results presented here, understanding how these pathways fit together and how and if these molecular pathways are influenced by neuronal activity will provide an integrated perspective on how vertebrate habituation learning occurs *in vivo*.

## Online Methods

### Zebrafish Husbandry, Mutagenesis, Transgenesis, and Genotyping

All animal protocols were approved by the University of Pennsylvania Institutional Animal Care and Use Committee (IACUC).

Mutant alleles were generated using CRISPR-Cas9 as previously described^32^. sgRNA were designed using ChopChop v2^76^, oligos were annealed and directly ligated into pDR274^77^. sgRNA were then synthesized using the T7 MEGAshortscript kit (Ambion) and purified using the MEGAclear kit (Ambion). Cas9 protein (PNA Bio) and sgRNAs were mixed and injected into 1-cell stage TLF embryos. G_0_ injected larvae were raised and outcrossed to identify and establish heterozygous carrier lines.

To generate *Tg*(*hsp70-hip14-p2a-mKate*), Gateway cloning was employed to combine *hip14-p2a-mKate* into a pDest vector containing the *hsp70* promoter and I-sce1 restriction sites. I-sceI transgenesis was performed as previously described^78^ in 1-cell stage embryos from *hip14+/-* incrosses. Transgenic lines were identified as previously described^33^. Briefly, 24 hpf F_1_ offspring of G_0_-injected fish were placed into Eppendorf tubes and incubated at 37°C for 30 min, recovered in petri dishes for 60 min, and screened for mKate with a fluorescent stereomicroscope (Olympus MVX10).

Fish carrying *Tg*(*GFFDMC130A*) and *Tg*(*GFF62A*) were provided by Dr. Koichi Kawakami^79,80^. *Tg*(*UAS:gap43-citrine*) fish were provided by Dr. Jonathan Raper^81^.

Adult zebrafish and embryos were raised at 29°C on a 14-h:10-h light:dark cycle. *hip14* and *kcna1a* mutations were genotyped using the KASP method with proprietary primer sequences (LGC Genomics). For genotyping in the context of *Tg*(*hsp-hip14-p2a-mKate*) we developed a dCAPS assay using the dCAPS program (http://helix.wustl.edu/dcaps/dcaps.html) to design appropriate primers, with the reverse primer binding in the intron adjacent to the mutation^82^.

### Behavior Testing

Behavioral experiments were performed on 4–5 dpf larvae and behavior was analyzed using FLOTE software as previously described^30,40^. Larvae were arrayed in a laser-cut 36-well acrylic dish attached via an aluminum rod to a vibrational exciter (4810; Brüel and Kjaer, Norcross, GA), which delivered acoustic vibrational stimuli (2ms duration, 1000 Hz waveforms). Acoustic startles were identified using defined kinematic parameters (latency, turn angle, duration and maximum angular velocity). Behavior was recorded at 1000 frames per second using a Photron UX50 camera suspended above the dish. Habituation was calculated as: % Habituation = (1-[response frequency **Stimuli 21-30**] ÷ [response frequency **baseline**])*100.

### WGS and Molecular Cloning of *hip14* and *kcna1a*

Mutant cloning was performed as previously described^29,32^. Briefly, pools of 50 behaviorally identified *p174 and p181* mutant larvae were made and genomic DNA (gDNA) libraries were created. gDNA was sequenced with 100-bp paired-end reads on the Illumina HiSeq 2000 platform, and homozygosity analysis was done using 463,379 SNP markers identified by sequencing gDNA from ENU-mutagenized TLF and WIK males as described previously^29^.

### Ca^+2^ Imaging

Combined Ca^+2^ imaging and acoustic startle experiments were performed as described previously^14^. Larvae were head-fixed in glass-bottomed petri dishes with 2% low-melt agarose diluted in bath solution containing 112mM NaCl, 5mM HEPES pH 7.5, 2mM CaCl_2_, 3mM glucose, 2mM KCl, 1mM MgCl_2_^83^. Tails were freed to permit behavioral responses to tap stimuli, and dishes were filled halfway with bath solution. 5-6 baseline responses were obtained by delivering acoustic tap stimuli every 2 minutes using a stage-mounted speaker. Short term habituation was then induced by delivering 25 tap stimuli with a 5 second interstimulus interval. Ca^+2^ transients were captured using a spinning disk confocal microscope, while behavioral responses were recorded at 500fps using a camera suspended above the larva. For lateral dendrite analysis during habituation, ΔF/F0 values were normalized to account for indicator sensitivity-related decreases in soma activity during action potentials as previously described^14^.

### Pharmacology

SC79 (Sigma #0749), U0126 (Cell Signaling Technology #9903S), and rolipram (Sigma #R6520) were dissolved in 100% DMSO and administered at a final concentration of 1% DMSO. All compounds were administered for 30 minutes prior to testing with the exception of SC79, which was administered chronically as previously described^29^ (replacing media daily from 3-5dpf).

### Electrophysiology in N2a cells

All recordings were performed at room temperature (20 - 21°C). Data were acquired with an Axopatch 200B amplifier at 5 kHz. Currents were filtered by an eight-pole Bessel filter at 1 kHz and sampled at 5 kHz with an 18-bit A/D converter. Electrode capacitance was compensated electronically, and 60% of series resistance was compensated with a lag of 10 μs. Electrodes were made from thick-walled PG10150-4 glass (World Precision Instruments). HEKA Pulse software (HEKA Eletronik, Germany) was used for data acquisition and stimulation protocols. Igor Pro was used for graphing and data analysis (WaveMetrics, Inc.). Leak subtractions were not applied in the current study.

The N2a mouse neuroblastoma cell line was cultured in Eagle’s minimum essential medium supplemented with 10% FBS and 0.5 × penicillin/streptomycin (Invitrogen) at 37°C in a humidified incubator with 5% CO_2_. Whole-cell currents were recorded in N2a cells 48 hours after transfection. The pipette solution contained (in mM): 140 K^+^, 1 Mg^2+^, 1 Ca^2+^, 30 Cl^-^, 11 EGTA and 10 Hepes, pH 7.3 adjusted by methanesulfonic acid, ∼300 mOsm. The bath solution contained (in mM): 150 Na^+^, 5.4 K^+^, 1.5 Ca^2+^, 1 Mg^2+^, 150 Cl^-^, 20 glucose, 1 μM TTX, and 10 Hepes, pH 7.4 adjusted by methanesulfonate, ∼320 mOsm. The electrode resistance was 1.5-3 MΩ in these recording conditions.

### ABE Palmitoylation Assays

HEK293 cells were transiently transfected with 1µg total of DNA using Mirus LT1 transfection reagent. The ABE assays were performed 15 hours after transfection as previously described^59^.

### Immunohistochemistry

Animals expressing UAS:Citrine under the control of the GFFDMC130a driver were fixed in 4% Paraformaldehyde and 0.25% Triton (PBT). Fish were stained as previously described^84^. First they were washed in PBT, incubated in 150 mM Tris-HCl, pH 9 for 5 minutes at room temperature, and then 15 minutes at 70°C, washed with PBT, permeabilized in 0.05% Trypsin-EDTA for 45 minutes on ice, washed with PBT, blocked with PBT + 1% bovine serum albumin + 2% normal goat serum + 1% DMSO, and then incubated for 48 hours (as previously described^61^) at 4°C with rocking in blocking solution containing antibodies against GFP (1:500 Chicken anti-GFP Aves Labs, #GFP-120) and Kv1.1 (1:200 Rabbit anti-Kv1.1; Sigma, #AB5174). Samples were then washed in PBT and incubated overnight at 4°C with rocking in secondary antibodies diluted in block solution. Larval jaws were dissected away, and fish mounted ventral side up in Vectashield. 1024×1024 resolution stacks were then collected with an inverted confocal microscope (Zeiss LSM880), using a 63x water immersion objective. Fiji was used to quantify Kv1.1 signal blind to genotype. Sum projections were generated for the confocal slices encompassing the Mauthner axon initial segment. Regions of interest were drawn over the Mauthner AIS segment from the medial extent of the soma to the point at which the axon begins its posterior turn, extending anteriorly and posteriorly to encompass a region of approximately the same width as the Mauthner soma. Total fluorescent intensity in the Kv1.1 channel within these regions of interest was then calculated and divided by the area of the ROI.

### Heat Shock-Induced *hip14* Rescue

To induce expression of Hip14-p2a-mKate, 4- or 5-dpf larvae were arrayed in individual wells of a 96- well plate and placed at 37°C for 40 min in a thermocycler. Larvae were then placed in Petri dishes and allowed to recover for 140 minutes in the 5dpf condition (or 1 day in the 4dpf condition). Importantly, the transgene had no effect on habituation learning under normal temperature raising conditions (28°C) indicating that *hip14* expression is dependent on heat-shock.

### Statistics

Statistical analyses, including calculation of means, SD and SE, were performed using Prism (GraphPad). Significance was assessed using t-tests or ANOVA with Tukey’s post-hoc test for multiple comparisons as indicated.

## Acknowledgements

The authors would like to thank Drs. Kawakami and Raper for the *Tg:GFFDMC130a* and *UAS- GAP43*-*Citrine* fish lines, respectively. The authors would also like to thank Dr. Andrea Stout and Jasmine Zhao at the Cell and Developmental Biology Microscopy Core who provided advice and assistance with image acquisition and analysis, and the University of Pennsylvania Next-Generation Sequencing Core. We also thank the Granato lab members as well as Drs. Marc Wolman, Roshan Jain, and Kurt Marsden for technical advice and feedback on the manuscript. This work was supported by grants to MG (NIH R01 MH109498), EW (NIH R01 CA181633 and ACS RSG-15-027-01), and JKF/ZM (NIH R01 DC012538). JCN was supported by a NRSA: 5F32MH107139. FC was supported by the Haverford Velay Scholars program. AF was supported by the Haverford KINSC Summer Scholars program.

## Author Contributions

JCN, EW, ZM, JKF, and MG designed the experiments; JCN, EW, FC, AF and ZM performed the experiments, JCN, EW, and ZM analyzed data; JCN and MG wrote the paper.

---

**Supplemental Figure 1.**
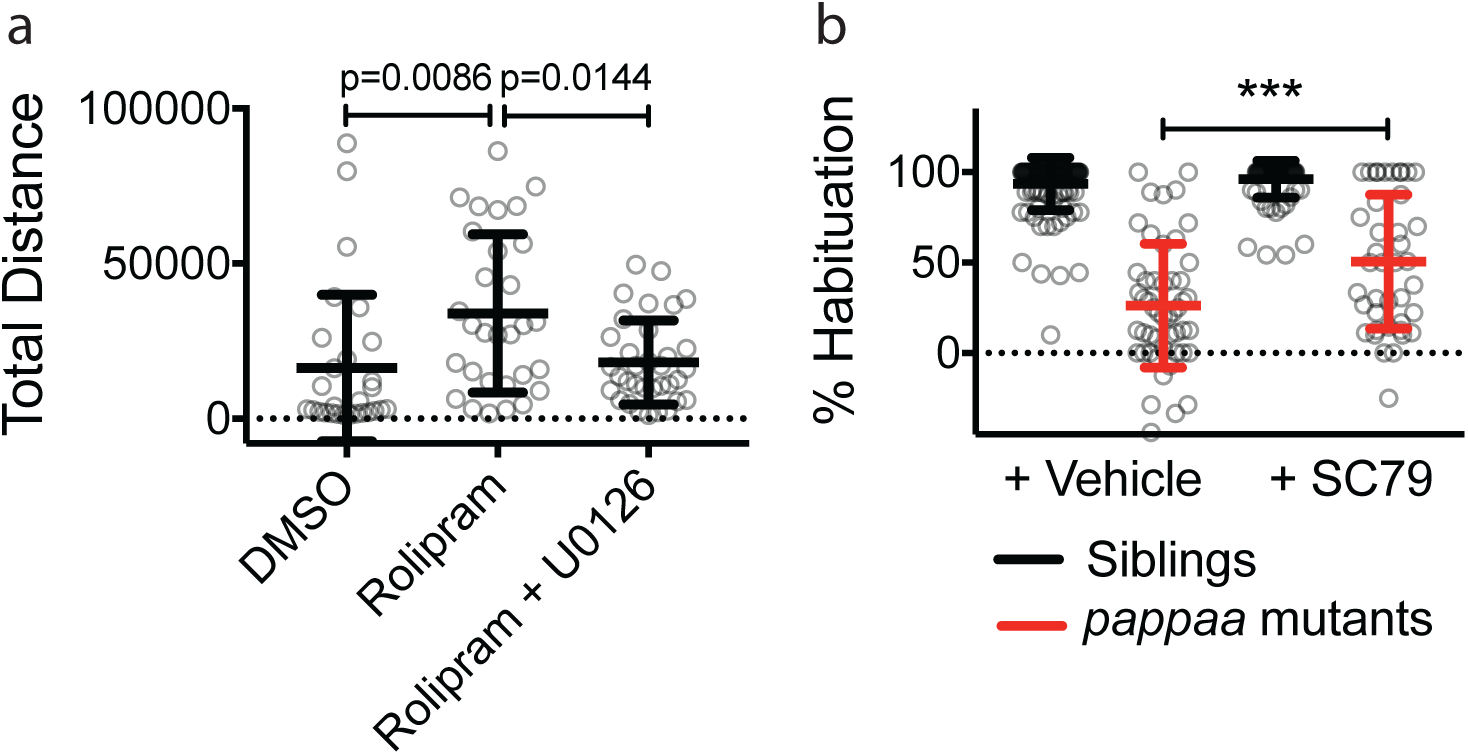
Drug Controls. (a) 10μM Rolipram applied 30 minutes prior to and throughout behavior testing as in **Fig. 3b** induces behavioral hyperactivity as previously described^85^, measured here as total distance traveled per larva in a 30 minute period, following 30 minutes of acclimation to behavior testing arena. MEK inhibitors are reported to restore normal activity levels^85^ and do so here in the case of 1μM U0126 (or DMSO control) applied with Rolipram 30 minutes prior to and throughout behavior testing. (b) 1μM SC-79 (or DMSO control) applied from 3dpf-5dpf and throughout behavior testing partially restores habituation learning in *papp-aa* mutants. *** = p<0.0001.

